# Transcriptomic profiling of mouse mammary tumors enables prognostic and predictive biomarker discovery for human breast cancers

**DOI:** 10.64898/2026.02.28.707759

**Authors:** Matthew D. Sutcliffe, Kevin R. Mott, Tulay Yilmaz-Swenson, Brooke M. Felsheim, Alexander V. Lobanov, Anna R. Michmerhuizen, Patrick D. Rädler, Denis O. Okumu, Xiaping He, Adam D. Pfefferle, Stephanie Dance-Barnes, Michael P. East, Daniel P. Hollern, Timothy C. Elston, Gary L. Johnson, Charles M. Perou

**Affiliations:** Lineberger Comprehensive Cancer Center, University of North Carolina at Chapel Hill, Chapel Hill, NC 27599, USA; Department of Genetics, School of Medicine, University of North Carolina at Chapel Hill, Chapel Hill, NC 27599, USA; Curriculum in Bioinformatics and Computational Biology, University of North Carolina at Chapel Hill, Chapel Hill, NC 27599, USA; Department of Biological Sciences, DePaul University, Chicago, IL 60604, USA; Department of Pharmacology, School of Medicine, University of North Carolina at Chapel Hill, Chapel Hill, NC 27599, USA; Salk Institute for Biological Studies, University of California, San Diego (UCSD), La Jolla, CA 92037

## Abstract

The development and validation of prognostic and predictive biomarkers in breast cancer is limited by the availability of well-annotated datasets linking tumor molecular features to treatment response and survival outcomes. To address this need, we generated an extensive mouse models dataset comprised of 26 immunocompetent mammary tumor models spanning diverse genetic backgrounds, epithelial–mesenchymal states, the basal–luminal axis, and distinct immune microenvironments. For each model, we measured survival under no treatment, immune checkpoint inhibition (ICI), and carboplatin/paclitaxel chemotherapy. We performed RNA-seq on baseline tumors and on 7-day on-treatment samples for both regimens.

Using baseline murine tumor gene expression features, we trained a machine learning Elastic Net model that predicted survival outcomes on multiple human breast cancer datasets with performance comparable to that of existing prognostic assays. We next trained models for ICI benefit, using either the untreated or 7-day ICI treated samples; both models predicted ICI benefit on human ICI treated datasets, with the 7-day treated tumor model showing better performance. We also developed a predictor of carboplatin/paclitaxel response that performed well in mice but did not generalize to human chemotherapy cohorts.

Finally, we compared multiple computational approaches, including XGBoost, random forests, and support vector regression; all methods successfully predicted survival outcomes, with Elastic Net offering the best performance and interpretability. These results indicate conserved cancer biology between mouse and human tumors for prognosis and ICI response and establish this large preclinical dataset with linked phenotypic and genomic data, as a resource for benchmarking computational methods for survival prediction.

**Significance:** The development of a genomically and phenotypically diverse murine tumor dataset with linked treatment outcomes establishes a robust translational resource to develop, test, and benchmark clinically relevant prognostic and therapeutic response biomarkers.

## INTRODUCTION

Breast cancer is a heterogeneous disease with different clinical and molecular subtypes, immune contexts, and responsiveness to treatment. Triple-negative breast cancer (TNBC) remains a particularly aggressive and clinically challenging subtype that shows a high degree of genetic and phenotypic heterogeneity^1–3^. For early-stage TNBC patients, immune checkpoint inhibition (ICI) with pembrolizumab and 4 chemotherapeutic agents is the standard of care^4,5^. However, variable response rates and survival outcomes to this immunotherapy and chemotherapy regimen are seen and reinforce the need for biologically informed biomarkers that can guide treatment decisions. In addition, human clinical cohorts typically suffer from the logistical difficulty of obtaining serial samples during treatment to investigate the molecular features of response and drug induced changes, and from confounding variables such as trial eligibility criteria that skew the biology of the patient set in known and unknown ways.

Preclinical models are an important resource for addressing some of the deficiencies and challenges present in human translational research. Patient-derived xenografts (PDXs) have widely been used because they preserve many human tumor features and enable drug screening of human tumors. While ongoing efforts are being made to reconstitute a more complete immune system in PDXs,^6^ or in organoid cultures,^7,8^ their lack of an intact adaptive immune system that can evolve remains a deficiency that can severely limit the ability to study drugs targeting adaptive immunity. In contrast, immunocompetent genetically engineered mouse models, including syngeneic transplant models,^9–13^ maintain a fully intact immune system and allow for therapeutic testing and mechanistic interrogation of tumor biology and response where the immune system is thought to be important.

Prior studies have shown that mouse mammary tumor models can replicate many of the gene expression programs and histological subtypes of human breast cancer^11,14,15^. This disease-state conservation suggests that machine learning algorithms trained on high-quality preclinical data could be used to predict human disease-state outcomes. Nevertheless, many existing mouse transcriptomic datasets lack on-treatment measurements, particularly with matched survival data, and almost always are limited to a handful of distinct preclinical models. Thus, there is a need for a comprehensive and well-annotated preclinical dataset that can connect murine tumor biology to human clinical response, and with sufficient sample size for de novo model building.

Here, we present a comprehensive transcriptomic and phenotypic murine dataset comprising 331 bulk RNA sequencing (RNA-seq) samples from 26 genetically diverse and immunocompetent mouse mammary tumor models profiled longitudinally and under two relevant treatment regimens. This dataset enables a direct linkage between tumor states and both off- and on-treatment outcomes. Additionally, we demonstrate that machine learning algorithms trained on the mouse dataset can predict both disease outcomes and response to immunotherapy on independent human cancer cohorts, with performance metrics comparable to established clinical assays. This dataset can serve as a powerful resource for additional biomarker discovery and/or validation, which can also enable further hypothesis generation in a testable system.

## METHODS

### Animal models

All genetically engineered mouse models have been previously published (see Table 1) except for the p53RB-426, p53RB-624, and p53RB-626 tumor models. These transplantable tumor lines were derived from 3 different spontaneous mammary tumors that developed in the mammary glands of K14-Cre (FVB-Tg(KRT14-cre)8Brn/Nci, RRID:MGI:3618476) Trp53^fl/fl^ (FVB.129P2-Trp53^tm1Brn^/Nci, RRID:MGI:3618506) Rb1^fl/fl^ (FVB;129-Rb1^tm2Brn^/Nci, RRID:MGI:3618505) triple transgenic females in an FVB background. Final triple transgenics were generated by initial breeding of Trp53^fl/fl^ mice with Rb1^fl/fl^ mice to obtain dual transgenic homozygotes, which were then crossed with homozygous K14-Cre carriers to obtain heterozygous triple transgenic offspring that was subsequently bred and propagated into homozygosity for the Trp53^fl/fl^ and Rb1^fl/fl^ transgenes. Mechanistically, in K14-Cre; Trp53^fl/fl^; Rb1^fl/fl^ mice, Cre recombinase expression is driven by a Keratin 14 (K14) promoter. In K14 expressing cells, Cre recombinase recognizes the loxP sites of the Trp53^fl/fl^ and Rb1^fl/fl^ alleles, leading to a recombination event that results in Trp53 and Rb1 null alleles. Single transgenics, Trp53^fl/fl^, Rb1^fl/fl^, and K14-Cre were obtained from the National Cancer Institute mouse repository (NCI repository strain numbers 01XC2, 01XC1, and 01XF1, respectively). The K14-TRT model^16^ is a cell line derived from a spontaneous K14-Cre Trp53^fl^/^fl^ Brca1^fl^/^fl^ mouse as described in Hollern et al.^17^

**Table 1:** Mouse tumor models used in this study.

| Tumor Model | Background | Genetics | Tumor Source | Subtype | Molecular Subtype | Reference |
| --- | --- | --- | --- | --- | --- | --- |
| 2153 | BALB/c | Trp53 −/− | Syngeneic Transplant | TNBC | Luminal-like | <sup>9</sup> |
| 2208 | BALB/c | Trp53 −/− | Syngeneic Transplant | TNBC | Luminal-like | <sup>9</sup> |
| 2224 | BALB/c | Trp53 −/− | Syngeneic Transplant | TNBC | Basal-like | <sup>9</sup> |
| 2225 | BALB/c | Trp53 −/− | Syngeneic Transplant | TNBC | Basal-like | <sup>9</sup> |
| 2250 | BALB/c | Trp53 −/− | Syngeneic Transplant | TNBC | Luminal-like | <sup>9</sup> |
| 2336 | BALB/c | Trp53 −/− | Syngeneic Transplant | TNBC | Basal-like | <sup>9</sup> |
| 9263 | BALB/c | Trp53 −/− | Syngeneic Transplant | TNBC | Basal-like | <sup>9</sup> |
| 4T1 | BALB/c | Trp53 mut, Pik3cg mut | Syngeneic Transplant | TNBC | Luminal-like/Claudin-low | <sup>70</sup> |
| E0771 | C57/BL6 | Trp53 mut | Syngeneic Transplant | TNBC | Luminal-like/Claudin-low | <sup>71</sup> |
| EGAN-ILC | C57/BL6 | Cdh1/Pik3ca mut | Autochthonous | TNBC | Normal-like | <sup>72</sup> |
| K14TRT | FVB | K14-Cre; Trp53 f/f Brca1 f/f | Syngeneic Transplant | TNBC | Claudin-low | <sup>16</sup> |
| KPB1-A | FVB | K14-Cre; Trp53 f/f Brca1 f/f | Syngeneic Transplant | TNBC | Basal-like/Claudin-low | <sup>13</sup> |
| KPB1-B | FVB | K14-Cre; Trp53 f/f Brca1 f/f | Syngeneic Transplant | TNBC | Basal-like | <sup>13</sup> |
| KPB25L | FVB | K14-Cre; Trp53 f/f Brca1 f/f | Syngeneic Transplant | TNBC | Basal-like | <sup>17</sup> |
| KPB25L-APOBEC | FVB | K14-Cre; Trp53 f/f Brca1 f/f; Apobec3 overexpression | Syngeneic Transplant | TNBC | Basal-like | <sup>17</sup> |
| KPB25L-UV | FVB | K14-Cre; Trp53 f/f Brca1 f/f; UV | Syngeneic Transplant | TNBC | Basal-like | <sup>17</sup> |
| p53RB-426 | FVB | K14-Cre; Trp53 f/f RB f/f | Syngeneic Transplant | TNBC | Claudin-low | This paper |
| p53RB-624 | FVB | K14-Cre; Trp53 f/f RB f/f | Syngeneic Transplant | TNBC | Claudin-low | This paper |
| p53RB-626 | FVB | K14-Cre; Trp53 f/f RB f/f | Syngeneic Transplant | TNBC | Claudin-low | This paper |
| T11 | BALB/c | Trp53 −/− | Syngeneic Transplant | TNBC | Claudin-low | <sup>10</sup> |
| T11-APOBEC | BALB/c | Trp53 −/−; Apobec3 overexpression | Syngeneic Transplant | TNBC | Claudin-low | <sup>17</sup> |
| T11-UV | BALB/c | Trp53 −/−; UV | Syngeneic Transplant | TNBC | Claudin-low | <sup>17</sup> |
| T12 | BALB/c | Trp53 −/− | Syngeneic Transplant | TNBC | Claudin-low | <sup>73</sup> |
| C3-TAG | FVB | SV40 T-antigen | Autochthonous | TNBC | Luminal-like | <sup>74</sup> |
| MMTV-PYMT | FVB | PyVT expression | Autochthonous | TNBC | Luminal-like | <sup>75</sup> |
| MMTV-Neu* | FVB | Her2/NEU expression | Autochthonous | HER2+/ER- | Luminal-like | <sup>76</sup> |

All animal work was conducted according to the Institutional Animal Care and Use Committee (IACUC) guidelines. All experiments were performed using female mice without blinding. For syngeneic transplant models, tumor propagation and transplantation was performed as previously described.^17,18^ Recipient mice were matured to 12 weeks of age and then injected with 100,000 to 500,000 cells depending upon the model. Following implantation, mice were monitored for tumor growth 2–3 times per week by caliper measurement. Once the tumor diameter reached 5 mm in any dimension, mice were randomized into treatment or control groups.

### Chemotherapy

Treated mice received carboplatin (50 mg/kg, Pfizer) and paclitaxel (10 mg/kg, Teva Pharmaceuticals) once per week by intraperitoneal injection. Mice were weighed twice weekly and monitored for adverse effects throughout treatment. Animals exhibiting signs of treatment-related toxicity were euthanized according to IACUC guidelines and removed from the study.

### Immunotherapy

Immunotherapy antibodies were administered intraperitoneally in 100 uL injection volumes twice per week. The following antibodies and doses were used: anti-PD1 (250 ug, Bio X Cell, BE0146, RRID:AB_10949053), anti-CTLA4 (125 ug, Bio X Cell, BE0164, RRID:AB_10949609), CD40AG (100 ug, Bio X Cell, BE0016-2, RRID:AB_1107647), IgG2a isotype control (125 ug, Bio X Cell, BE0089, RRID:AB_1107769), and IgG2b isotype control (125 ug, Bio X Cell, BE0086, RRID:AB_1107791). For the chemotherapy experiments, we also performed controls, which did not receive any IgG2a isotype control treatments. We have compared the survival of IgG2a controls to no isotype antibody controls and observed no statistically significant survival differences; one set of controls was used for the survival ratio calculations for on-treatment survival times for both the chemotherapy and immunotherapy treatments.

### Survival study

Baseline body weight was recorded prior to therapy initiation. Tumor size was measured by caliper instrumentation and body condition score (BCS) was assessed every 2–3 days. Mice were humanely euthanized when tumors reached 20 mm in any dimension, or when multiple tumors were noted. Independent of tumor size, animals were also euthanized if body weight loss reached 20% or more of baseline or if BCS declined to 2 or below. These criteria were applied uniformly across all treatment groups and any of these events was considered an event for our definition of “endpoint-free survival”.

### Mouse tumor bulk RNA-seq

mRNAseq libraries were generated from up to 1 µg of total RNA using a TruSeq Stranded mRNA kit (Illumina 20020594) following the manufacturer’s protocol (Illumina 1000000040498). Paired-end sequencing was performed on either the Illumina HiSeq 2500 or Illumina NovaSeq 6000. FASTQ files were merged when necessary and aligned to the mm10 reference genome with Gencode M25 annotations^19^ using STAR 2.7.11b,^20^ and quantification was performed using Salmon 1.10.3^21^. Next, Salmon abundance estimates were upper quartile-normalized (i.e. 75^th^ percentile of non-zero values was scaled to 1000) and then log2-tranformed with the addition of 1 pseudocount. For differential expression analyses, Salmon abundance estimates were rounded to the nearest integer prior to analysis with DESeq2^22^.

### Gene expression modules

894 published gene expression signatures representing cell types, biological functions, and cell signaling pathways were previously identified based on published gene lists. 840 signatures were based on the per-sample median expression of a defined set of genes, 44 “special models” were calculated as the correlation to a pre-defined centroid, and another 10 “special models” are algorithms that have previously been published^23–31^. 3 of these modules were the basis for clinical prognostic signatures (Research Prosigna – “ROR_P_Model_JCO.2009_PMID.19204204”, Research OncotypeDX – “GHI_RS_Model_NEJM.2004_PMID.15591335”, and Research Mammaprint – “Pcorr_NKI70_Good_Correlation_Nature.2002_PMID.11823860”) and an additional 3 prognostic modules were previously published (“TNBC_Clinically_Relevant_Good.26_BCR.2011_PMID.21978456”, “TNBC_Clinically_Relevant_Poor.230_BCR.2011_PMID.21978456”, and “TNBC_Clinically_Relevant_Poor.26_BCR.2011_PMID.21978456”) and thus these features were excluded from all Elastic Net and other machine learning prognostic model building.^28,29,32,33^ A list of these modules and their associated Entrez IDs genes are listed in Supplementary Table 3.

To compute gene expression module scores for each dataset, gene symbols or Entrez IDs were first mapped to Ensembl IDs. If present, duplicate Ensembl IDs were collapsed by summing counts. Upper quartile-normalized counts were then log2-transformed with a pseudocount of 1. To calculate median-based modules, genes with less than 70% zero values were removed. Next, rows (genes) were median-centered, and then columns (samples) were column-standardized. For each median-based module, module scores were calculated as the per-sample median expression value of the intersecting gene list. To calculate centroid-based modules, rows (genes) were first standardized. Depending on the module, the parametric Pearson correlation, non-parametric Spearman correlation, or Euclidean distance from each sample to the predefined centroid was computed using the intersecting gene lists. Special algorithm modules were computed using a best approximation based on the original methods^23–31^. An R package to calculate all 894 gene expression module scores is available at GitHub: https://github.com/perou-lab/Breast-Cancer-Modules

### Human gene expression sets

The Swedish Cancerome Analysis Network – Breast (SCAN-B) is a multicenter prospective center study including clinical and RNA-sequencing data^34^. Clinical data for 6,329 samples were obtained from a previous publication^35^. FASTQ files from 6,329 samples were obtained from collaborators at Lund University as previously described^36^. The UNC-337 DNA-microarray dataset (GEO: GSE18229) is a curated dataset of 337 distinct tumors with clinical follow-up that was previously published^30^. For the NKI-295 dataset, which contains 295 distinct tumors, the gene expression and clinical data were obtained from a previous publication^37^; the NKI-295 microarray log10 ratios were obtained from the “SeventyGeneData” R package^38^. Probes with > 30% missing values were removed. Probes were then mapped to gene symbols. Remaining missing values were imputed using the “impute” R package with k-nearest neighbors of k = 10. Duplicate genes were collapsed using the mean. Log10 ratios were converted to log2 ratios.

For the Bassez et al. (2021) single-cell RNAseq dataset, read count and phenotypic data was obtained from the original manuscript^39^. For each patient sample, counts from single cells were aggregated via summation to generate pseudobulk samples prior to normalization. For the Gide et al. (2019) melanoma dataset, FASTQ files were obtained from the European Nucleotide Archive^40^ (ENA) under accession number PRJEB23709. Paired clinical data was obtained from the original manuscript^41^. The CALGB 40603 clinical trial data was obtained from the NCBI database of Genotypes and Phenotypes (dbGaP, RRID:SCR_002709, RNA-seq data: phs001863.v1.p1, phenotypic data: phs003801.v1.p1), as previously described^42,43^. For all human RNA-seq data, FASTQ files were aligned to the hg38 reference genome with Gencode (RRID:SCR_014966) v36 annotations (Gide melanoma) or v47 annotations (SCAN-B) using STAR (RRID:SCR_004463) and quantified with Salmon. Salmon abundance estimates were upper quartile-normalized and log2-transformed with a pseudocount of 1.

### Machine learning models

All machine learning models employed a bootstrap resampling strategy with 100 iterations. At each bootstrap iteration, mouse tumor models were sampled with replacement to create training sets of the same size (i.e. number of tumor models). Out-of-bag (OOB) sets consisted of tumor models not selected in the training set and therefore varied in size at each iteration. In all cases, biological replicates from each tumor model were restricted to either the training set or the OOB set to prevent overestimation of performance.

### Elastic net regression

Elastic net models were implemented using the “glmnet” R package. A two-dimensional hyperparameter grid was explored with alpha values ranging from 0 to 1 in increments of 0.1 (mixing parameter between ridge and LASSO regularization), and lambda values from 10^-4 to 10^4 on a logarithmic scale with increments of 0.1 log units. Models were trained to predict either the log-transformed median survival time for untreated models or the log-transformed survival time ratio (treated time / untreated time) for treatment response models. For each (alpha, lambda) combination, models were fit on training data and evaluated on both training and OOB sets. Performance metrics included mean squared error (MSE) and concordance index (c-index) for survival prediction. Hyperparameter values were chosen based on the minimum MSE in the OOB sets.

### Random forest

Random forest models were implemented using the “ranger” R package. Hyperparameter grids included: number of trees (500, 1000, 1500), features per split (mtry: 10, 20, 40, 80), minimum node size (5, 10, 15, 20), maximum tree depth (5, 10, 15, 20), and sample fraction (0.5, 0.632, 0.7). Models were trained to predict log-transformed median survival time. Performance was evaluated using root mean squared error (RMSE) and c-index on training and OOB sets.

### XGBoost

XGBoost models were implemented using the “xgboost” R package. Hyperparameter grids included: learning rate (eta: 10^-2 to 10^-0.5), maximum tree depth (1, 2, 3), minimum split loss (gamma: 0, 0.1, 0.5, 1), L2 regularization (lambda: 10^-1 to 10^2), L1 regularization (alpha: 10^-1 to 10^2), row sampling fraction (0.5, 0.8, 1.0), and column sampling fraction (0.1, 0.2, 0.5). Models were trained for 100 rounds. Performance was evaluated using RMSE and c-index on training and OOB sets.

### Support vector regression

Support vector regression (SVR) models were implemented using the “e1071” R package with epsilon-insensitive loss. Hyperparameter grids included: kernel type (linear, radial basis function, polynomial), regularization parameter C (0.01, 0.1, 1, 10, 100), epsilon (0.01, 0.05, 0.1, 0.2), kernel coefficient gamma (0.001, 0.01, 0.1, 1), and polynomial degree (2, 3, 4) for polynomial kernels. Performance was evaluated using RMSE and c-index on training and OOB sets.

### Immunotherapy-based gene expression analyses

For immunotherapy response analysis, differential expression between anti-PD1 + anti-CTLA4 treated and untreated samples was performed using DESeq2^22^. Only samples from tumor models exhibiting a response to anti-PD1 + anti-CTLA4 (survival time ratio > 1.25, “responders”) were considered for differential expression analysis. A design matrix accounting for treatment and tumor model effects was used: design = ∼ Treatment + Tumor Model. Differential expression results were filtered for statistical significance (p-value < 0.01) and effect direction (log2 fold change > 0 for upregulated genes).

Linear regression was performed to identify genes associated with response to immunotherapy (as defined by the median survival time ratio) in untreated tumors. For each gene, a linear regression model was fit: log(treated time / untreated time) ∼ expression. Genes with positive coefficients and p-value < 0.01 were considered biomarker candidates.

Genes were identified as biomarkers if they met the criteria for both the differential expression analysis and the correlation analysis. Additionally, unnamed mouse genes and retained intron genes were excluded from the final list. Gene set enrichment analysis (GSEA) was performed on biomarker gene lists using the Molecular Signatures Database (MSigDB, RRID:SCR_016863) Hallmark gene sets^44,45^. Enrichment results were filtered by false discovery rate (FDR) q-value < 0.05.

### Data availability

The underlying mouse tumor gene expression is available at GSE223630, GSE124821, and GSE304115. See Supplementary Table 1 for GEO sample metadata. Mouse survival data can be found in Supplementary Table 2. Elastic Net models with weights and immune response signature genes can be found in Supplementary Table 4.

### Code availability

The code to calculate the 894 gene expression modules is available as an R package at GitHub: https://github.com/perou-lab/Breast-Cancer-Modules. The code used to generate the machine learning models for this study is available at https://github.com/perou-lab/mouse-mammary-tumor-modeling.

## RESULTS

### Mouse tumor model cohort, transcriptomic profiling, and baseline characterization

To establish a comprehensive resource for developing and testing gene expression-based prognostic and predictive computational models, we performed RNA-seq on fresh frozen tumors from 26 genetically distinct mouse mammary cancer models (Table 1); of these models, 22 are syngeneic transplants, and 4 are autochthonous models. These mouse tumor models are all immunocompetent and include 25 TNBC models and 1 HER2+ model, yet represent a diversity of known genomic subtypes, including luminal, basal, and claudin-low^11–13^. These models also represent the most common somatic mutations in TNBC, including functional loss of TP53 and/or BRCA1, and/or RB1^46^. For each mouse tumor model, we collected bulk RNA-seq data from biological replicates of untreated tumors (n = 3–9 each) and separately monitored tumor growth over time using multiple untreated tumors (n ≥ 7 each) (Figure 1A). From the tumor growth monitoring study, we defined survival time as the time in days from an initial tumor diameter of 0.5 cm to an IACUC-mandated endpoint (including tumor greater than 2 cm, multiple tumors, tumor ulceration, poor overall animal health). Median survival times ranged widely across tumor models, from 9 days for p53Rb-426 to 43 days for KPB25L-APOBEC, representing substantial diversity in baseline tumor progression rates (Figure 1B).

**Figure 1:**
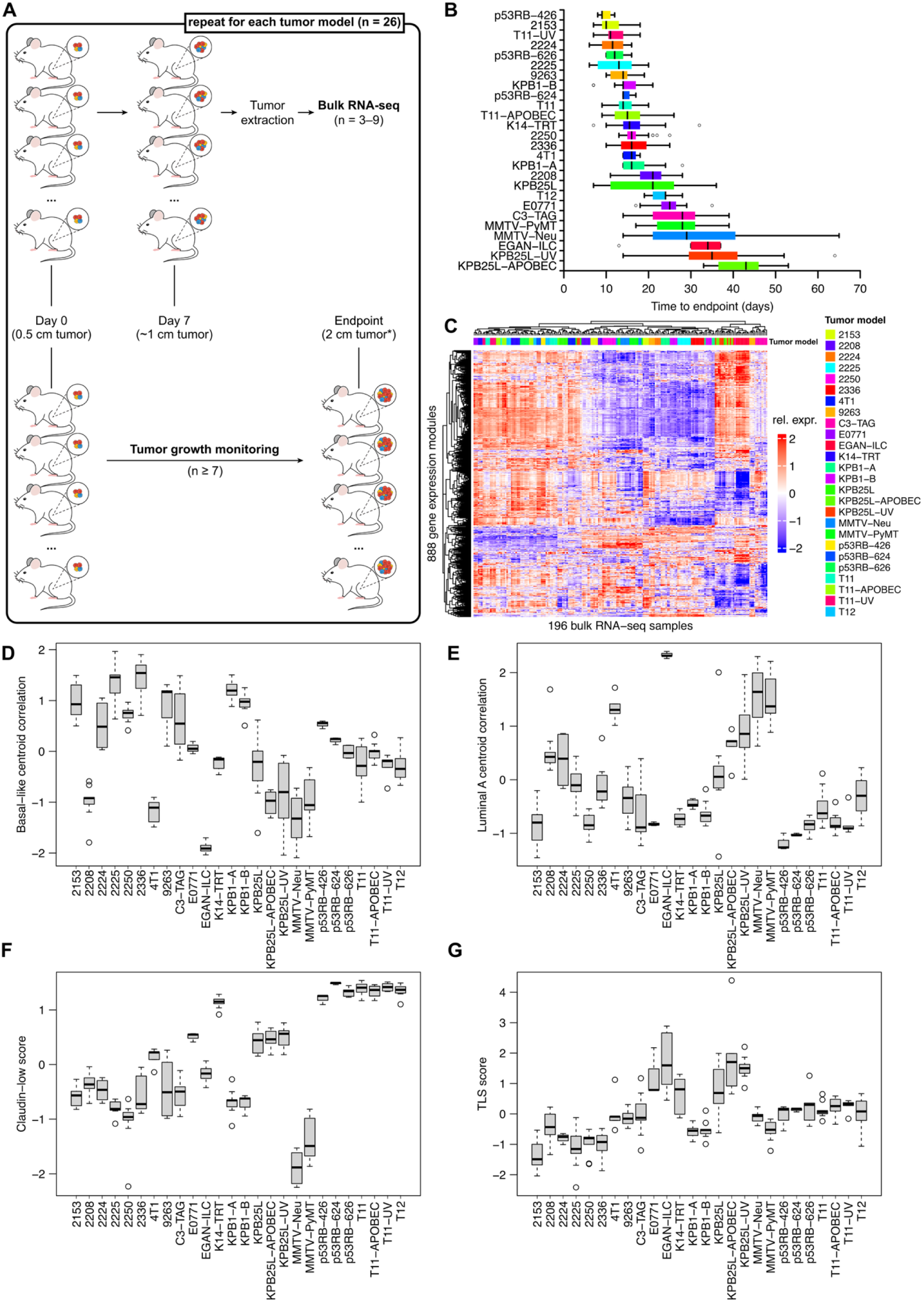
Transcriptomic and phenotypic profiling of 26 mouse mammary tumor models. (A) Summary of untreated bulk RNA-sequencing samples and survival endpoint measurements for each tumor model. 3–9 biological replicates were collected and processed for bulk RNA-sequencing. (B) Boxplots of time-to-endpoint measurements for 26 mouse mammary tumor models. Center line represents median, boxes represent the 25^th^ to 75^th^ percentile, and whiskers represent minimum or maximum values within 1.5 times the interquartile range from the 25^th^ or 75^th^ percentile. (C) Heatmap of 196 untreated bulk RNA-seq samples across 26 tumor models (columns). Rows represent row-standardized scores for literature-derived gene expression modules (see Methods). (D) Correlation of each sample to the basal-like subtype centroid. (E) Correlation of each sample to the luminal-like subtype centroid. (F) Inverse distance of each sample to the Claudin-low centroid. (G) Median module score for Tertiary Lymphoid Structures (TLS).

To improve interpretability and reduce dimensionality of the bulk RNA-seq data, we summarized the gene expression features into a curated set of 840 median expression gene signatures, 44 centroid-based models, and 10 special algorithms, which in total represent a diversity of key biological pathways, cellular states, and cell types (see Methods), similar to what is encompassed within Molecular Signatures Database^44^. These gene expression “modules” have been extensively published by our laboratory,^43,47–49^ and were compiled from independent publications, the Molecular Signatures Database (MSigDB) Hallmark gene sets,^50^ and from literature relevant to cancer biology and the tumor microenvironment. We then used these 888 genomic features (excluding 6 known prognostic features) in a hierarchical clustering analysis of 196 untreated murine tumor RNA-seq samples (Figure 1C). In almost all instances, the mouse model biological replicates clustered together, with mixing between tumor models only seen between a parental line and its mutagenized derivative (e.g. KPB25L and KPB25L-UV).

In breast cancer, a tumor’s genomic subtype is a well-studied feature. Based upon previous publications,^11,17,18^ we assigned a molecular “intrinsic” subtype to each of the 26 models (Table 1). Subtype mapping using selected gene expression modules revealed alignment of mouse tumor models to human breast cancer molecular subtypes. Basal-like centroid correlations were predominantly highest among the p53-null models, including 2336, 2225, and 2153 (Figure 1D), whereas luminal A centroid correlations were highest in the EGAN-ILC, MMTV-Neu, and MMTV-PyMT luminal models (Figure 1E). In contrast, several models were highly enriched for the claudin-low mesenchymal phenotype, including the previously described T11, T12, and newly created p53^fl/fl^ Rb^fl/fl^ models (Figure 1F). Finally, a gene expression signature representing tertiary lymphoid structures (TLS) scores, thus representing adaptive immunity, also varied across models, with higher TLS genomic scores indicating an immune-rich microenvironment (Figure 1G). Together, these data establish a transcriptomic landscape capturing the basal–luminal axis, claudin-low/mesenchymal states, and the immune microenvironment across the mouse tumor model cohort.

### Development of a mouse-derived Elastic Net prognostic model with cross-species validation

To evaluate the prognostic model-building capability of the mouse tumor dataset, we trained an Elastic Net regression model using 888 gene expression features (after removing 6 known prognostic features) and the log2 of the tumor model-specific median survival time as the “clinical” outcome for each untreated sample. We applied bootstrap resampling with out-of-bag (OOB) evaluation across tumor models to identify optimal values for the alpha (ridge/LASSO mixing term) and lambda (regularization weight) hyperparameters (Figure 2A). In each bootstrap simulation, all tumor biological replicates per model were restricted to either the training set or the OOB set to prevent overfitting (see Methods). Following hyperparameter selection of alpha = 0.1 and lambda = 0.126, we then re-fit the final model on the entire mouse dataset. The final model identified a set of 106 features of varying weights (40 most highly weighted features are shown in Figure 2B) representing a diverse set of biological features. Top weighted features included basal (poor outcome) or luminal (good outcome) gene expression modules, as well as immune signatures (good outcome). We then applied the final model to the median expression profile of the untreated mouse tumors and stratified endpoint-free survival into tertiles, with a concordance index (c-index) of 0.939 indicating strong agreement between predicted and observed survival (Figure 2C).

**Figure 2:**
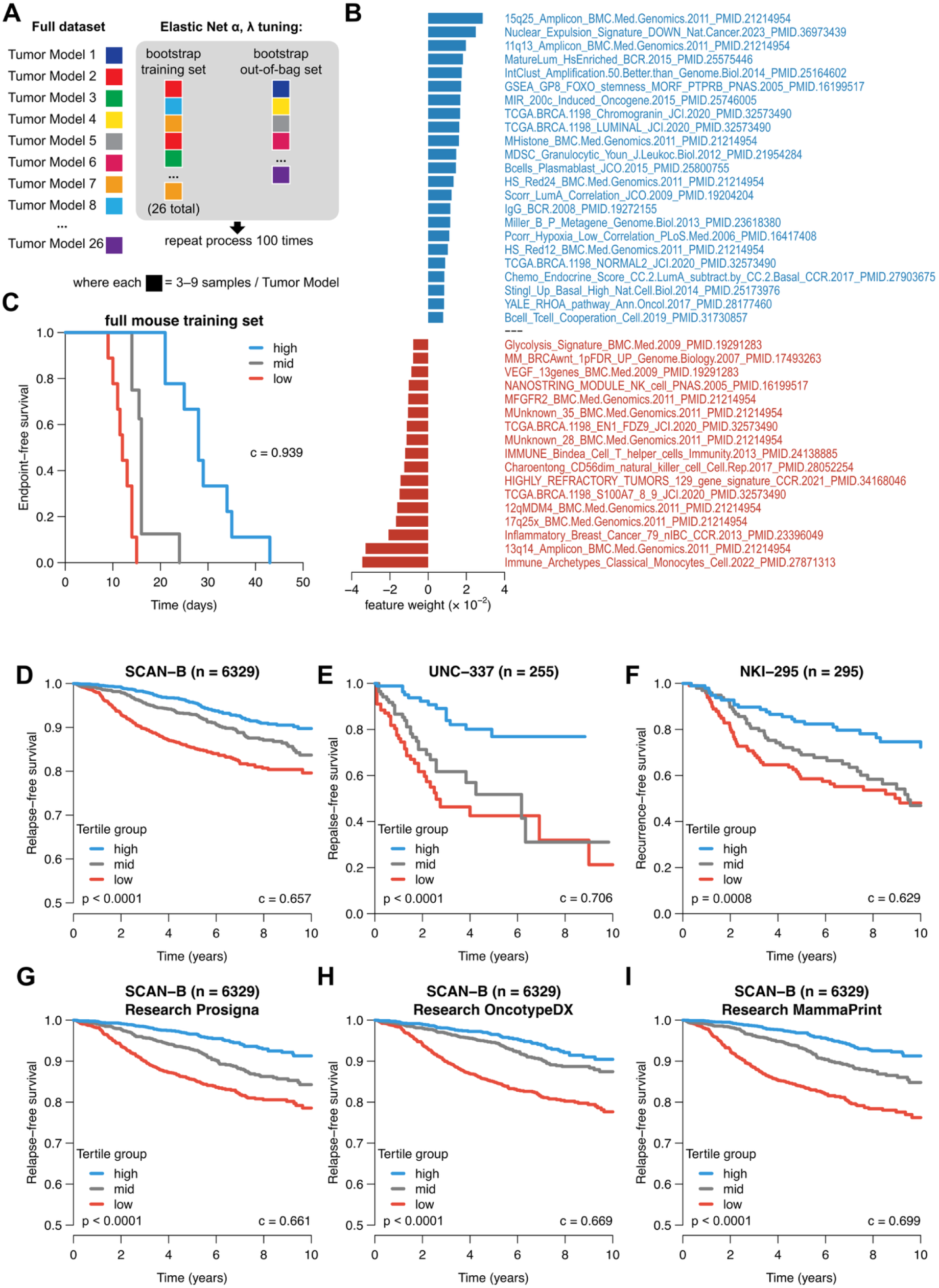
Elastic Net prognostic modeling of untreated mouse tumors and application to human datasets. (A) Bootstrapping approach to select optimal combination of alpha and lambda (see Methods). (B) Top 40 features from final fitted Elastic Net model. Higher expression of features with positive weights (blue) predicts longer survival times and higher expression of features with negative weights (red) predicts shorter survival times. (C) Mouse training set model performance, with predicted scores split into equal tertiles. (D) Mouse-derived Elastic Net model applied to the human SCAN-B dataset. Predictions were split into tertiles prior to plotting. Tertile p-values by log-rank test are shown. (E) Mouse-derived Elastic Net model applied to the human UNC-337 microarray dataset. Predictions were split into tertiles prior to plotting. Tertile p-values by log-rank test are shown. (F) Mouse-derived Elastic Net model applied to the human NKI-295 microarray dataset. Predictions were split into tertiles prior to plotting. Tertile p-values by log-rank test are shown. (G) Research version of Prosigna PAM50 ROR-P prognostic model applied to the human SCAN-B dataset. Predictions were split into tertiles prior to plotting. Tertile p-values by log-rank test are shown. (H) Research version of OncotypeDX prognostic model applied to the human SCAN-B dataset. Predictions were split into tertiles prior to plotting. Tertile p-values by log-rank test are shown. (I) Research version of MammaPrint prognostic model applied to the human SCAN-B dataset. Predictions were split into tertiles prior to plotting. Tertile p-values by log-rank test are shown.

Next, we assessed the translational performance of this mouse-derived Elastic Net prognostic model on three different human datasets. We calculated the 888 gene expression module scores on three human breast cancer datasets, including SCAN-B^34,35^ (RNA-seq based), UNC-337^30^ (DNA microarray), and NKI-295^33,51^ (DNA microarray). In SCAN-B, the model successfully stratified according to relapse-free survival (RFS) with an overall performance c-index of 0.657 (Figure 2D). Similar performance was observed in the UNC-337 cohort (c-index = 0.706, Figure 2E) and the NKI-295 cohort (c-index = 0.629, Figure 2F), noting that the NKI cohort is similar to our murine training set in that these patients received no systemic adjuvant therapies^33,51^. For comparison, we benchmarked the mouse-derived risk predictor against established clinical prognostic assays by applying analogous tertile-based stratifications in SCAN-B using research versions of Prosigna^28^, OncotypeDX^29^, and Mammaprint^33^ (Figures 2G–I). Interestingly, the human-derived prognostic models (c-index values ranging from 0.661 – 0.699) showed similar performance to that of the mouse-derived Elastic Net model, supporting the utility of this mouse dataset in identifying clinically-relevant genomic features and might serve as a biomarker. Lastly, we examined within-clinical subtype performance (i.e., ER+/HER2-, HER2+, and TNBC), and we observed that the murine model performs similarly well in HER2+ and ER+/HER2- subsets, but not on TNBC, again reflecting the performance of the human prognostic models (Figure S1).

### Predictive modeling of response to immune checkpoint inhibition

Building upon the prognostic modeling framework, we aimed to build a predictive model for survival on immune checkpoint inhibition. To do this, we tested 23 of the mouse tumor models (excluding EGAN-ILC, K14-TRT, and KPB25L-APOBEC) with combined anti-PD1 + anti-CTLA4 therapy (ICI), which is used on human melanoma patients and currently being tested on TNBC patients^52,53^. In parallel to the untreated data, we collected at least 3 biological replicates per model, following 7 days of ICI treatment for bulk RNA-seq processing. In addition, we evaluated endpoint-free survival times on continuous ICI treatment (administered twice weekly). To quantify the response to ICI therapy, we defined the median survival time ratio in treated vs. untreated mice for each model (Figure 3A). While many of the tumor models did not benefit from ICI treatment, several tumor models were exquisitely sensitive to ICI therapy, including E0771, T11-APOBEC, and T11-UV. Using the same set of 888 gene expression modules described earlier, we fit an Elastic Net regression model on the 7-day treated murine tumors to predict the survival time ratio and identified 87 selected features (25 most highly weighted features are shown in Figure 3B). Among the features with positive weights (i.e. higher expression predicting sensitivity to ICI) were many T-cell signatures, with the top weighted feature being a T-regulatory feature, which is the target of ICI.

**Figure 3:**
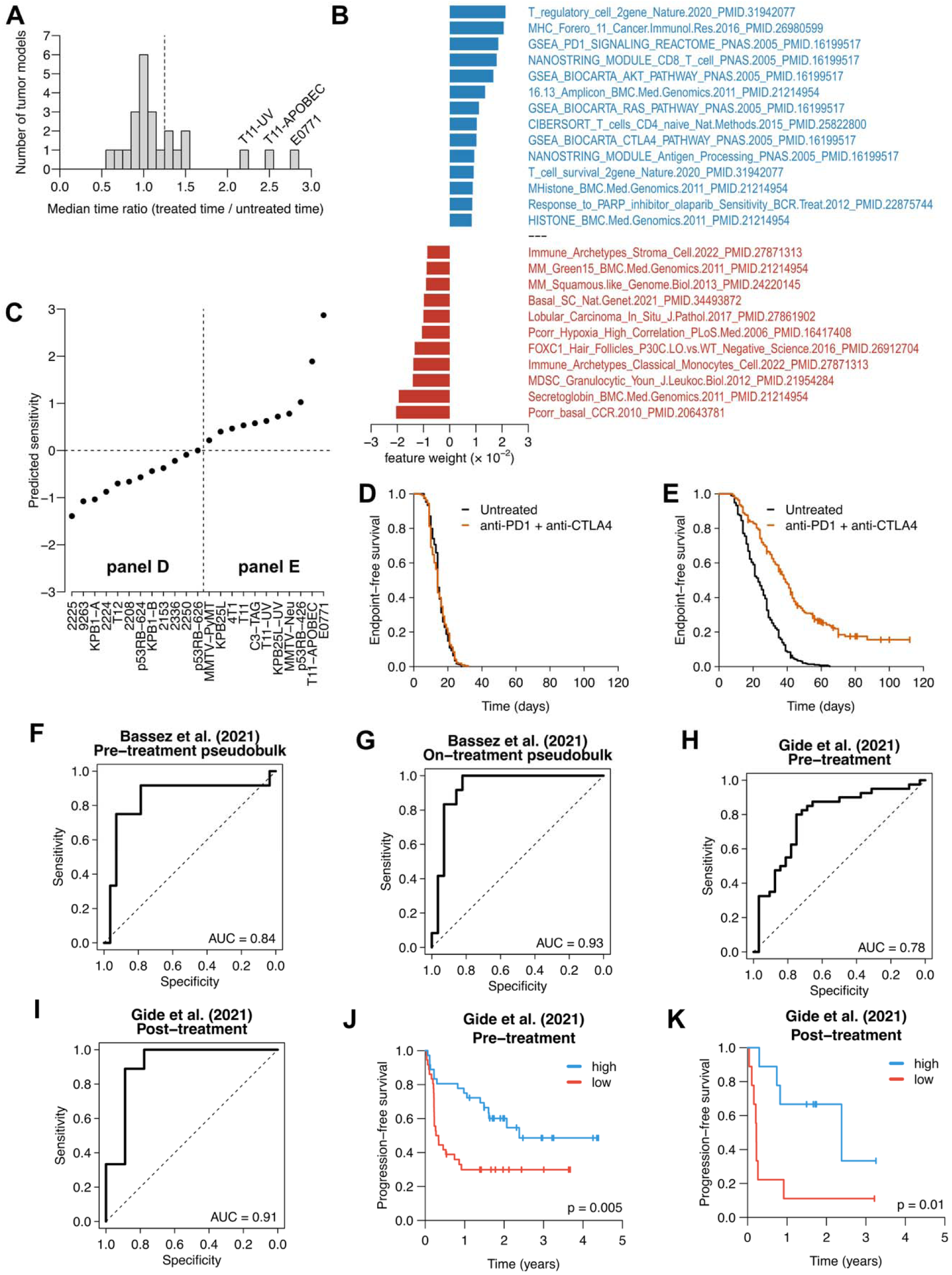
Elastic Net predictive modeling of ICI response in untreated mouse tumors and validation in human datasets. (A) Histogram of mouse tumor model response to ICI. A time ratio of 1 indicates no difference in median survival time in ICI-treated vs. untreated tumors. (B) Top 25 weighted features in final Elastic Net ICI response predictor. Positive weights (blue) predict sensitivity to ICI and negative weights (red) predict tumor resistance. (C) Elastic Net model predictions for the median expression profiles of untreated mouse tumors. (D) Pooled ICI- and untreated-tumor survival data for mouse tumor models with below-median predicted sensitivity. (E) Pooled ICI- and untreated-tumor survival data for mouse tumor models with above-median predicted sensitivity. (F) Receiver operating characteristic (ROC) curve of Elastic Net model predictions for Bassez pseudobulk samples, used to predict T-cell expansion in pre-treatment samples. (G) ROC curve of Elastic Net model predictions for Bassez pseudobulk samples, used to predict T-cell expansion in on-treatment samples. (H) ROC curve of Elastic Net model predictions for Gide samples, used to predict tumor response (partial/complete response vs. stable/progressive disease) using the pre-treatment samples. (I) Receiver operating characteristic (ROC) curve of Elastic Net model predictions for Gide samples, used to predict tumor response (partial/complete response vs. stable/progressive disease) using the post-treatment samples. (J) Kaplan-Meier plots for Elastic Net model predictions for Gide pre-treatment samples, with predictions split into above- and below-median groups. (K) Kaplan-Meier plots for Elastic Net model predictions for Gide post-treatment samples, with predictions split into above- and below-median groups.

We applied this Elastic Net model onto the untreated mouse tumors to evaluate pre-treatment biomarker activity, using the median expression profiles of the tumor biological replicates as input to rank-order the sensitivity predictions (Figure 3C). When the underlying survival data was divided into those tumor models with below-median predicted sensitivities and above-median predicted sensitivities (Figure 3D, E), we observed strong predictive capabilities. To test this Elastic Net model on external data, we used two human datasets. The first set is Bassez et al. (2021)^39^, which is a pre- and on- anti-PD1 treatment single-cell RNA-seq dataset for which the outcome of interest was T-cell expansion (as measured by TCR-sequencing) following pembrolizumab treatment in 40 breast cancer patients. We computed gene expression module scores using the pseudobulk counts of each scRNA-seq human sample. We then separately tested the murine ICI Elastic Net model on the pre- and on-treatment samples. We observed strong predictive capability for T-cell expansion endpoint in both the pre-treatment (Figure 3F) and on-treatment (Figure 3G) samples. Next, we tested the Gide et al. (2019)^41^ dataset, in which melanoma patients received anti-PD1 monotherapy or anti-PD1 + anti-CTLA4 combination therapy. Similarly, we observed strong predictive capability for tumor response as defined by RECIST criteria (complete or partial response vs. stable or progressive disease) using both pre-treatment (n = 72, Figure 3H) and post-treatment (n = 18, Figure 3I) bulk RNA-seq samples. Additionally, the Elastic Net model successfully predicted progression-free survival (PFS) using both pre-treatment (Figure 3J) and post-treatment (Figure 3K) samples.

Finally, we also constructed an Elastic Net model using the untreated mouse tumor bulk RNA-seq samples instead of the 7-day ICI-treated samples. Here, the model selected fewer immune signatures (Figure S2A), yet like the previous analysis, we observed good stratification of mouse tumors based on pre-treatment median gene expression (Figure S2B–D). This model successfully predicted T-cell expansion on the Bassez breast tumor dataset (Figure S2E, F) but was less robust in predicting melanoma ICI response (Figure S2G–J), suggesting models derived from on-treatment samples may be more powerful, even when applied to pre-treatment tumors.

### ICI treatment biological discovery

Because our ICI sensitivity Elastic Net prediction model is based upon previously published gene expression modules as features, we sought to perform exploratory analyses using the original (individual gene-level) bulk RNA-seq data to identify finer grained biological features. We first divided the mouse tumor models into two groups, namely ICI “responders” and “non-responders” based on the survival time ratios (Figure 3A, dotted line). We first identified genes that were differentially expressed in the responders in the untreated vs 7-day treated tumors. In addition, we fit a linear regression model on the untreated tumors to identify a “sensitivity coefficient” for each gene. When combining the differential expression analysis with the sensitivity coefficient, we identified a set of 246 genes that both increased following ICI treatment in the responders and were individually predictive of sensitivity to ICI (Figure 4A). This gene list included many T-cell (e.g. Cd3[d, e, g], Cd8) and B-cell (e.g. Cd22, Cd40) genes, and gene set enrichment analyses identified immune-related HALLMARK gene sets, including INTERFERON_GAMMA_RESPONSE and ALLOGRAFT_REJECTION as the most highly enriched gene sets (Figure 4B).

**Figure 4:**
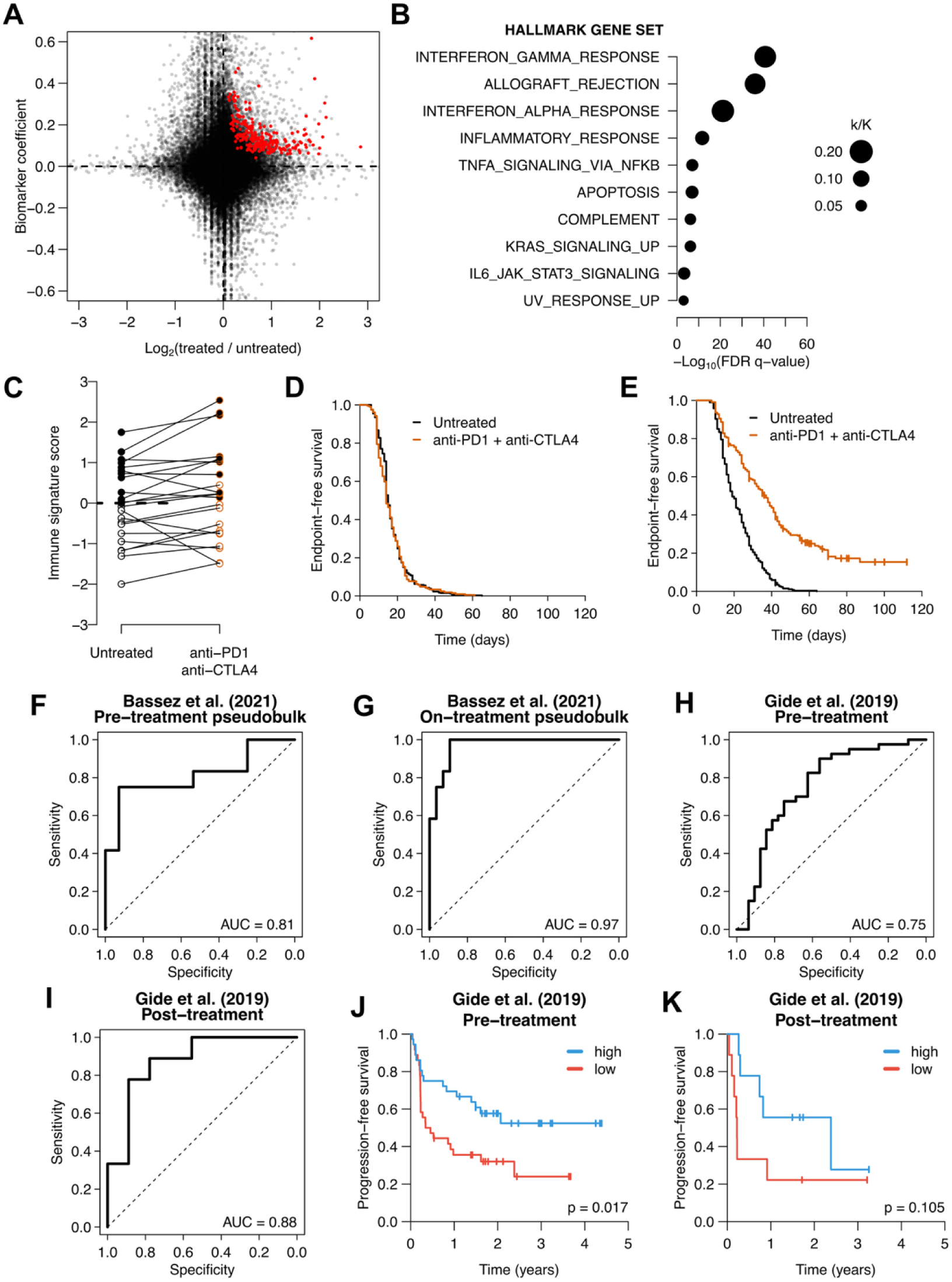
Agnostic ICI biomarker identification in ICI-treated mouse tumors. (A) Scatter plot of gene-based differential expression and biomarker analyses. X-axis represents fold change in ICI-treated vs. untreated mouse tumors categorized as responders. Y-axis represents the slope of the best fit line when median survival time ratio is predicted using untreated expression levels. 246 genes that were significantly increased in ICI-treated tumors and significantly correlated with survival outcomes are highlighted in red. (B) Gene Set Enrichment Analysis of 246 significant genes from (A). Top 10 HALLMARK pathways are shown. Size of circles represent the fraction of overlapping genes (k) over the size of the HALLMARK gene set (K). (C) Immune signature score calculated for median expression profiles of untreated and ICI-treated tumor models. Tumor models with above-median immune signature scores are filled in black. Tumor models with below-median immune signature scores are filled in white. (D) Pooled ICI- and untreated-tumor survival data for mouse tumor models with below-median immune signature scores. (E) Pooled ICI- and untreated-tumor survival data for mouse tumor models with above-median immune signature scores. (F) ROC curve for Bassez pseudobulk samples, where immune signature scores are used to predict T-cell expansion in pre-treatment samples. (G) ROC curve for Bassez pseudobulk samples, where immune signature scores are used to predict T-cell expansion in on-treatment samples. (H) ROC curve for Gide samples, where immune signature scores are used to predict tumor response (partial/complete response vs. stable/progressive disease) using the pre-treatment samples. (I) ROC curve for Gide samples, where immune signature scores are used to predict tumor response (partial/complete response vs. stable/progressive disease) using the post-treatment samples. (J) Kaplan-Meier plots for immune signature scores for Gide pre-treatment samples, with predictions split into above- and below-median groups. (K) Kaplan-Meier plots for immune signature scores for Gide post-treatment samples, with predictions split into above- and below-median groups.

We next created a new gene expression signature based upon this set of 246 genes, with the per-sample median expression as the “immune signature” score. We calculated immune signature scores in the median expression untreated and ICI-treated murine samples and observed an increase following treatment in many of the tumor models (Figure 4C), as would be expected. Further, we also considered this immune signature score as a biomarker of ICI treatment and split the mouse models into above-median and below-median expressing tumors before treatment. Here, we observe that those with below-median immune signature scores include the tumor models that are resistant to ICI (Figure 4D), while those with above-median immune signature scores displayed sensitivity to ICI (Figure 4E). Notably, compared to the Elastic Net model described previously, which has 87 modules based on 3020 unique genes, this much smaller list is similarly capable of stratifying mouse tumors by ICI response. To further evaluate this signature’s performance, we again utilized Bassez et al. 2021 and Gide et al. 2019 as test sets (Figures 4F–K) and observed qualitatively similar results to those of the ICI treated Elastic Net model.

Given the translatability of these murine computational models onto human tumor datasets, we next searched the 246 genes list for possible drug targets and observed that both the CD40 receptor and CD40 ligand were among the 246 genes in the immune signature. The CD40 receptor is a central regulator of immune response, primarily expressed on antigen-presenting cells (APCs), including dendritic cells and B cells. T cells can produce CD40 ligand and thus engage and activate CD40+ cells, leading to activated dendritic cells and B cells^54–56^. CD40 ligand, often referred to as CD40AG, is a monoclonal antibody therapy that is being tested on multiple human tumors including pancreatic tumors and melanoma^57–59^. Therefore, we tested a murine compatible version of a CD40AG antibody on ICI sensitive and resistant models, alone or in combination with ICI. We first tested an ICI-sensitive model (E0771), and observed that CD40AG alone, and in combination with anti-PD1, extended survival and exhibited similar benefit to the anti-PD1 + anti-CTLA4 combination (Figure 5A). Next, we tested the anti-PD1 + CD40AG combination on five p53-null tumor models that were resistant to the anti-PD1 + anti-CTLA4 combination. We found that two tumor models significantly benefited from anti-PD1 + CD40AG therapy (2208 (Figure 5B) and 2224 (Figure 5C)) and three did not (2153, 2336, and 2225, data not shown).

**Figure 5:**
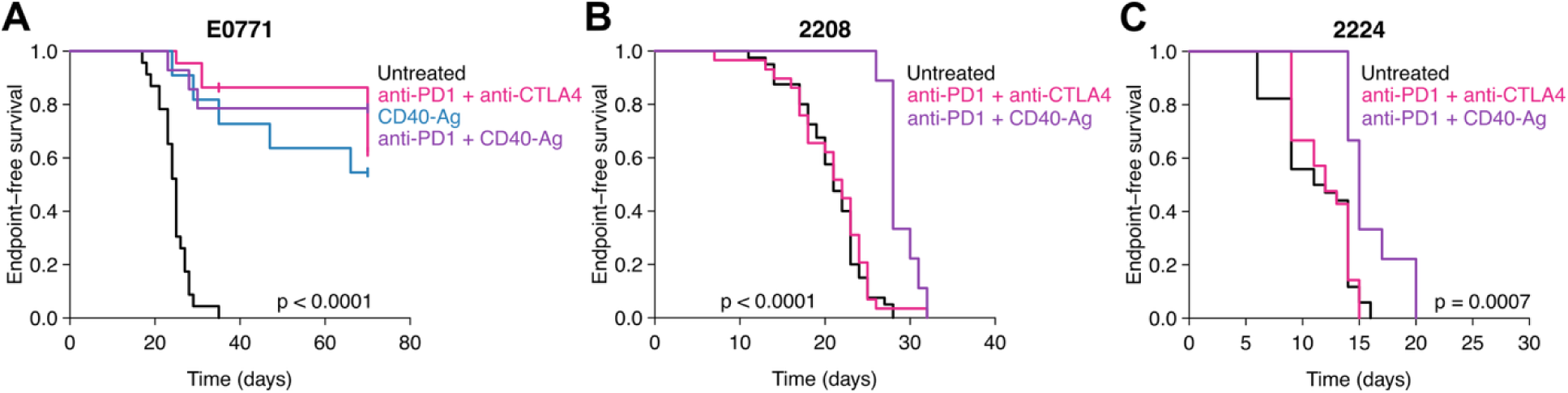
CD40 agonist treatment in ICI-sensitive and ICI-resistant tumor models. (A) Kaplan-Meier plots for E0771. P-value by log-rank test. (B) Kaplan-Meier plots for 2208. P-value by log-rank test. (C) Kaplan-Meier plots for 2224. P-value by log-rank test.

### Predictive modeling of response to carboplatin/paclitaxel chemotherapy

Because most early-stage TNBC patients receive multi-agent neoadjuvant chemotherapy as part of standard of care treatments, we were interested to see if the mouse tumor panel could be used to derive a chemotherapy benefit predictor. As seen with untreated and ICI treated mouse models, we observed varying sensitivities to carboplatin/paclitaxel treatment across our 24 models tested (Figure S3A), noting the three of the four most sensitive models were all derived from our Trp53^fl/fl^/Brca1^fl/fl^ model^13^. Next, we trained an Elastic Net model to predict carboplatin/paclitaxel treatment survival benefit using 7-day treated mouse tumor bulk RNA-seq samples and treated median survival time ratios. Using the same Elastic Net modeling approach, the resulting model identified several PARP inhibitor sensitivity modules among the most highly weighted features whose high expression predicted response (Figure S3B). Notably, when the Elastic Net model is applied to the median expression profiles of the untreated mouse tumors, all BRCA-deficient tumor models (KPB1-A, KPB1-B, KPB25L, KPB25L-UV) were predicted to have above-average sensitivity to carboplatin/paclitaxel (Figure S3C). Additionally, when the underlying survival data was split into above- and below-median sensitivity predictions, the Elastic Net model successfully predicts carboplatin/paclitaxel response compared to untreated (Figure S3D, E). However, when applied to human chemotherapy treated datasets, the Elastic Net model failed to robustly predict response or survival. From the SCAN-B cohort, we subset to the chemotherapy treated patients and evaluated predicted chemotherapy sensitivities; the Elastic Net had a modest prognostic effect in the SCAN-B chemotherapy subset (c-index = 0.58, p = 0.0009, Figure S3F); however, the high-score patients had worse outcomes. Additionally, when the Elastic Net model is applied to samples from CALGB 40603,^42^ a TNBC phase II trial testing the addition of carboplatin or bevacizumab to anthracycline- and taxane-based chemotherapy, the model failed to predict event-free survival or pathological complete response (Figure S3G, H).

### Comparative analysis of machine learning approaches

To assess whether alternative machine learning algorithms might improve computational model performance in our untreated mouse prognostic modeling, we trained XGBoost^60^, random forest, and support vector regression models using the murine dataset and testing on a human dataset. Similar to the Elastic Net regression training, we used bootstrapping to optimize hyperparameter selection (see Methods for details). With each of these approaches, we fit a model using the same mouse untreated tumor expression data and untreated endpoint-free survival times and evaluated these additional models on the full SCAN-B cohort (Table 2). All methods performed similarly, with Elastic Net achieving the best concordance index on the entire SCAN-B cohort. When analyzing clinical subtype-specific performance, similar patterns were observed with consistent performance for all methods on the ER+/HER2- and HER2+ subsets, and all showed reduced performance in TNBC. Given these similar performance metrics, we favor the Elastic Net modeling method due to its improved interpretability of selected features.

**Table 2:** Machine learning methods comparison for untreated prognostic model trained on mouse tumors and tested on human SCAN-B.

| **SCAN-B set** | **Elastic Net regression** | **XGBoost** | **Random Forest** | **Support Vector Regression** |
| --- | --- | --- | --- | --- |
| All | 0.657 | 0.619 | 0.633 | 0.651 |
| HR+/HER2- | 0.614 | 0.607 | 0.622 | 0.602 |
| HER2+ | 0.667 | 0.605 | 0.611 | 0.631 |
| TNBC | 0.530 | 0.571 | 0.566 | 0.526 |

## DISCUSSION

In this study, we created and used a phenotypically and genetically diverse panel of 26 mouse models of mammary cancer to develop prognostic and predictive transcriptomic-based machine learning models of survival and therapeutic response. We found that this large and diverse mouse tumor models data set captured much of the molecular heterogeneity of human breast cancers, particularly the spectrum ranging from differentiated luminal phenotypes to the more aggressive and less differentiated basal-like and claudin-low phenotypes. The conservation of the basal–luminal axis in murine models is known^11,61^ and well represented in this cohort, which allows for the interrogation of these biological programs and new hypotheses in a controlled, immunocompetent setting that may better allow for the translation of some findings from mouse models into human clinical breast cancer research. We believe a key advance here was to use the mouse in a manner more similar to what is done in human studies; namely, to base findings upon a phenotypically diverse population of “individuals”, instead of relying on one or two individuals and then extrapolating to a larger population. In addition, this murine dataset provides a unique resource often only seen with cell lines, where multiple treatments are performed on the same model, and on 10s of models, thus providing multiple individually determined treatment responses on the same model.

To efficiently characterize the high-dimensional bulk tumor RNA-seq data generated here, we employed a curated set of 894 gene expression modules; we now provide a R package that can run these 840 median gene signatures, 44 centroid-based modules, and 10 special algorithms (https://github.com/perou-lab/Breast-Cancer-Modules). This approach addresses a common challenge in cross-species genomics: the noise associated with individual genes and the precise linking of gene-by-gene orthologs. By summarizing the median expression of functionally related and/or co-expressed genes into modules, and by using previously rigorously identified special algorithms (i.e., OncotypeDX, MammaPrint, Prosigna), this analysis emphasizes biological processes and buffers the noise coming from the expression of individual genes. We have previously used these gene expression modules to identify prognostic and predictive features for early-stage HER2+ and TNBC patients,^48,49,43,42,62^ and here we show that they can be used in mouse datasets to predict murine model survival times and murine model response to immunotherapy. Unlike dataset-specific principal component analysis or metagene analysis, in which clusters of genes are used as input features in machine learning analyses, the gene sets presented here are independent of the original datasets and thus much more readily used across a variety of technologies, as indicated by our RNA-seq models working well on decades old DNA microarray data.

Using these gene expression features, we evaluated multiple machine learning approaches and determined that Elastic Net regression had comparable performance to more complex algorithms, including XGBoost^60^, random forest^63^, and support vector regression^64^. This finding is consistent with recent analyses evaluating risk of recurrence models in hormone receptor-positive breast cancer samples^65^. This is also consistent with similar studies using DNA methylation-based cancer features, in which Elastic Net was compared to decision-tree and support vector-based methods and found to be superior^66^. Further, the Elastic Net regression approach selects a subset of all features input and returns easily interpretable feature weights that explain the relative contribution and the directionality of each selected feature. Although importance metrics exist for tree-based algorithms, the relationships between features and predicted outcomes are non-linear and not necessarily monotonic, requiring cut points that vary greatly between decision trees.

Using the Elastic Net approach, we first developed a survival model entirely trained on our murine tumor dataset of 26 models represented by 196 RNA-seq samples, and it achieved performance comparable to established clinical tools (research versions) tested in the laboratory setting, including Prosigna, OncotypeDX, and MammaPrint, when evaluated on the same publicly available patient datasets (SCAN-B, NKI-295, UNC-337). While another prognostic predictor in breast cancer is not clinically needed, our results do further inform tumor biology and suggest existing prognostic models may be improved by including new immune system features. Importantly, these findings support the idea that the underlying tumor biology in our syngeneic transplant and autochthonous mouse models is conserved across species and that we have captured much of the molecular diversity of human breast cancer in our 26 mouse models dataset.

One of the more striking findings in this study was the success of our mouse model-derived ICI predictor, which worked well in two different human datasets spanning two different tumor types and “rediscovered” the ICI drug target (i.e., T-regulatory cells) as the top weighted Elastic Net model feature. In addition, when we performed more conventional exploratory analyses to identify treatment-induced features associated with response to ICI, we identified a rich set of adaptive immunity genes that performed well as a biomarker of response on human ICI-treated datasets, even when using just 246 genes. Given its simplicity as a single feature median-based signature, further testing through retrospective analysis of existing ICI randomized trials in TNBC might provide support for clinical implementation. In addition, this gene set includes many genes of mechanistic importance, including IFNγ and IL21. Using one of these 26 models (KPB25L-UV), we previously showed that administering neutralizing antibodies against IL21 at the same time as anti-PD1 + anti-CTLA4 antibodies inhibited the responsiveness of this model to ICI.^17^ Another mechanistically and possibly therapeutically important finding from this analysis was the identification of the CD40–CD40 ligand axis, which is a known immune cell crosstalk network often resulting in dendritic cell and/or B cell activation^67^. Therefore, we performed functional experiments using CD40AG antibodies in multiple different mouse models and saw single agent activity in one of our ICI responsive models (E0771), and activity with anti-PD1 in 2 out of 5 ICI-resistant models tested (Figure 5). These promising preclinical model results suggest testing CD40AG antibodies in TNBC patients either alone or in combination with anti-PD1 antibodies.

Given the success of our murine derived prognostic and ICI-specific predictor, it was surprising to determine that a similarly trained carboplatin/paclitaxel chemotherapy predictor largely failed when applied to human data (although it did perform well on the mouse training dataset). There are several possible explanations for this result, primarily that for human TNBC, there is no clinically used biomarker to predict chemotherapy response; thus, it clearly is quite challenging to identify such a biomarker using any model system. A second explanation for this failure, and the failure to have an established human TNBC chemotherapy biomarker, is that TNBC patients typically receive 3 or more chemotherapeutics in the early-stage setting, each with a distinct mechanisms of action, which we recapitulated here on our mouse models by giving carboplatin (DNA damage) and paclitaxel (microtubule inhibitor). In human TNBC, it is even more complex due to the inclusion of additional chemotherapeutics (i.e. cyclophosphamide and doxorubicin)^68^. Given this complexity of drugs, and its failure to generalize to human datasets, we would not advocate clinical testing of our murine-derived chemotherapy predictor. Finally, we did not perform murine experiments of chemotherapy plus ICI and an attempt to build a predictor of this, which is the treatment regimen used for human TNBC patients (i.e. KEYNOTE-522 regimen^4^), because we did not want to preface an ICI predictor on the foundation of a failed chemotherapy predictor.

Our study has several limitations, including that the biological replicate tumor samples used for bulk RNA-seq were different from those used for the survival studies. Intratumoral heterogeneity and stochastic biological variation between tumor replicates could introduce noise that impacts the predictive accuracy of our models. However, through hierarchical clustering of the bulk RNA-seq data, we showed that the transcriptomic states of mouse tumor biological replicates are stable and reproducible for the models we tested. Additionally, we mitigated this limitation by using tumor model- and treatment-specific median survival times when modeling prognosis and response to treatment to reduce the impact of individual mouse variability. A second limitation is that we separately tested ICI and chemotherapy, whereas in clinical settings, most TNBC patients receive both chemotherapy and ICI. A third limitation is that our mouse models set did not contain any Estrogen Receptor-positive, hormonally driven models, which is the most common clinical human breast cancer subtype; however, our mouse models did contain “luminal” murine tumors, and the basal-luminal axis was prognostic and translated to prognostic differences across human tumors. These results demonstrate that many clinically relevant features are conserved across species, while some are not, and thus while the mouse may not accurately model all aspects of human breast cancers, it does well modeling some, including baseline prognosis and many aspects of adaptive immunity^69^.

In summary, our diverse and “population-based” mouse mammary tumor dataset has both prognostic and predictive capabilities and extends these capabilities to human datasets. Additionally, the gene expression module features utilized in this study are publicly available and conveniently accessible as an R package. These modules can readily be extended with new or custom gene expression signatures and applied to new datasets to summarize and interpret the biology of mouse mammary or human breast cancer datasets. This murine data resource is a rich dataset for new discoveries and validation of existing biomarkers and should continue to interface with similar human datasets to test new hypotheses and computational methods.

## Author Contributions

M.D.S. conducted all computational analyses, developed the R package, and wrote the manuscript. K.R.M. and D.P.H. performed the mouse experiments and edited the manuscript. T.YS., A.D.P., and X.H. processed RNA-seq samples and edited the manuscript. B.M.F., A.V.L., A.R.M., and P.D.R. assisted with R package development and manuscript editing; A.V.L. and A.R.M. additionally processed and analyzed human datasets. S.DB. characterized the K14-TRT mouse model. D.O.O., M.P.E., T.C.E., and G.L.J. provided scientific input. C.M.P. conceptualized and supervised the project, provided overall support for all studies, and edited the manuscript. All authors contributed to manuscript writing and editing.

## Acknowledgments

Research was in part supported by funds from the NCI U01CA238475, Breast SPORE program (P50-CA058223), the Breast Cancer Research Foundation (BCRF-23-127), the Susan G. Komen (SAC-160074), National Institute of General Medical Sciences (5T32 GM067553), UNC iTOP (T32-CA24415), and the UNC Lineberger Triple Negative Breast Cancer Center. We would also like to thank the UNC LCCC Office of Genomics Research, the UNC LCCC Translational Genomics Lab (TGL) (RRID SCR_025231), and the UNC High-Throughput Sequencing Facility.

## Conflicts of Interest

C.M.P is an equity stockholder and consultant of BioClassifier LLC; C.M.P is also listed as an inventor on patent applications for the Breast PAM50 Subtyping assay.

