## Supplementary Figures and Tables for "Transcriptomic profiling of mouse mammary tumors enables prognostic and predictive biomarker discovery for human breast cancers": Supplementary_Figure_Descriptions.pdf

### Figure S1: Clinical subtype-specific performance of Elastic Net model and clinical tools on SCAN-B dataset.

(Rows) Elastic Net model, and research versions of prognostic tools.

(Columns) Clinical subtypes of breast cancer in the SCAN-B dataset.

**Figure S2: Elastic Net model of ICI response trained on untreated mouse tumors.**

(A) Top 25 weighted features in final Elastic Net ICI response predictor. Positive weights (blue) predict sensitivity to ICI and negative weights (red) predict tumor resistance.
(B) Elastic Net model predictions for the median expression profiles of untreated mouse tumors.
(C) Pooled ICI- and untreated-tumor survival data for mouse tumor models with below-median predicted sensitivity.
(D) Pooled ICI- and untreated-tumor survival data for mouse tumor models with above-median predicted sensitivity.
(E) Receiver operating characteristic (ROC) curve of Elastic Net model predictions for Bassez pseudobulk samples, used to predict T-cell expansion in pre-treatment samples.
(F) ROC curve of Elastic Net model predictions for Bassez pseudobulk samples, used to predict T-cell expansion in on-treatment samples.
(G) ROC curve of Elastic Net model predictions for Gide samples, used to predict tumor response (partial/complete response vs. stable/progressive disease) using the pre-treatment samples.
(H) Receiver operating characteristic (ROC) curve of Elastic Net model predictions for Gide samples, used to predict tumor response (partial/complete response vs. stable/progressive disease) using the post-treatment samples.
(I) Kaplan-Meier plots for Elastic Net model predictions for Gide pre-treatment samples, with predictions split into above- and below-median groups.
(J) Kaplan-Meier plots for Elastic Net model predictions for Gide post-treatment samples, with predictions split into above- and below-median groups.

**Figure S3: Elastic Net model of chemotherapy response.**

(A) Histogram of mouse tumor model response to carboplatin/paclitaxel. A time ratio of 1 indicates no difference in median survival time in chemotherapy-treated vs. untreated tumors.
(B) Top 25 weighted features in final Elastic Net chemotherapy response predictor. Positive weights (blue) predict sensitivity to chemotherapy and negative weights (red) predict tumor resistance.
(C) Elastic Net model predictions for the median expression profiles of untreated mouse tumors.
(D) Pooled chemotherapy- and untreated-tumor survival data for mouse tumor models with below-median predicted sensitivity.
(E) Pooled chemotherapy- and untreated-tumor survival data for mouse tumor models with above-median predicted sensitivity.
(F) Kaplan-Meier plots for Elastic Net model predictions in SCAN-B patients receiving chemotherapy, with predictions split into equal tertiles.
(G) Kaplan-Meier plots for Elastic Net model predictions in CALGB-40603 patients receiving chemotherapy, with predictions split into equal tertiles.
(H) Receiver operating characteristic (ROC) curve of Elastic Net model predictions of tumor response for CALGB-40603 patient samples.
