## Supplementary figures and images for "Transcriptomic profiling of mouse mammary tumors enables prognostic and predictive biomarker discovery for human breast cancers"

### Sutcliffe_et_al_SUPPLEMENTARY_FIGURES.pdf

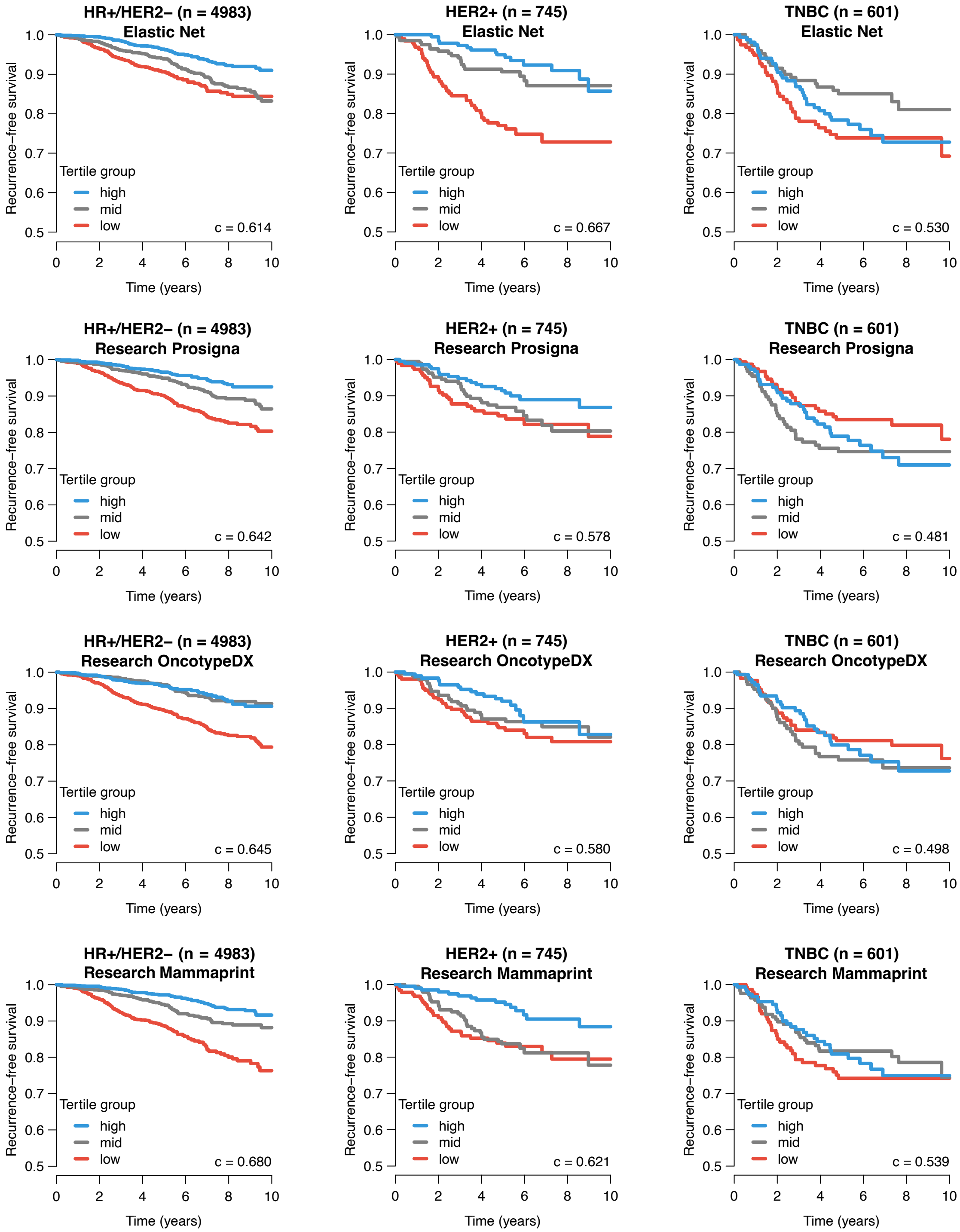

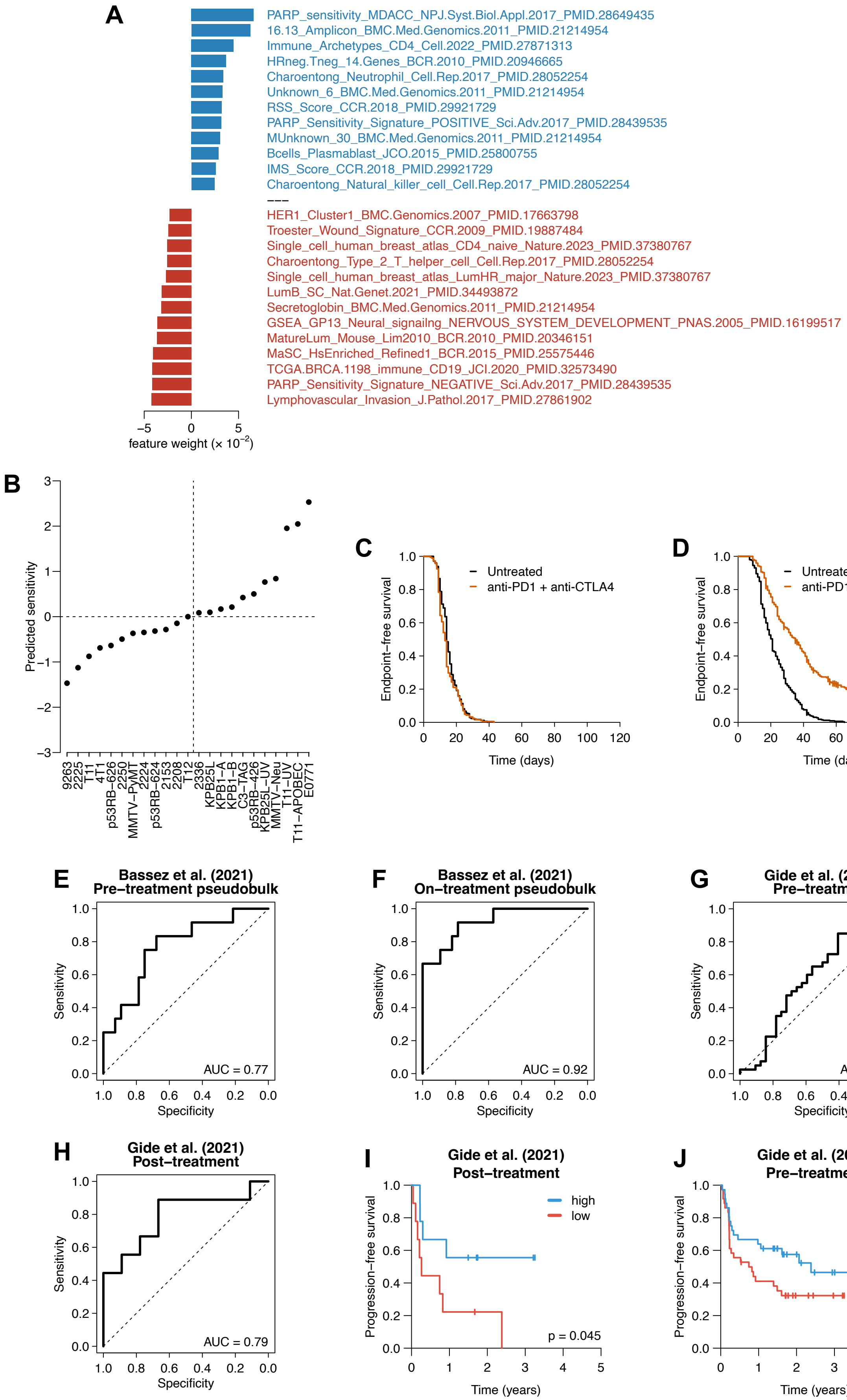

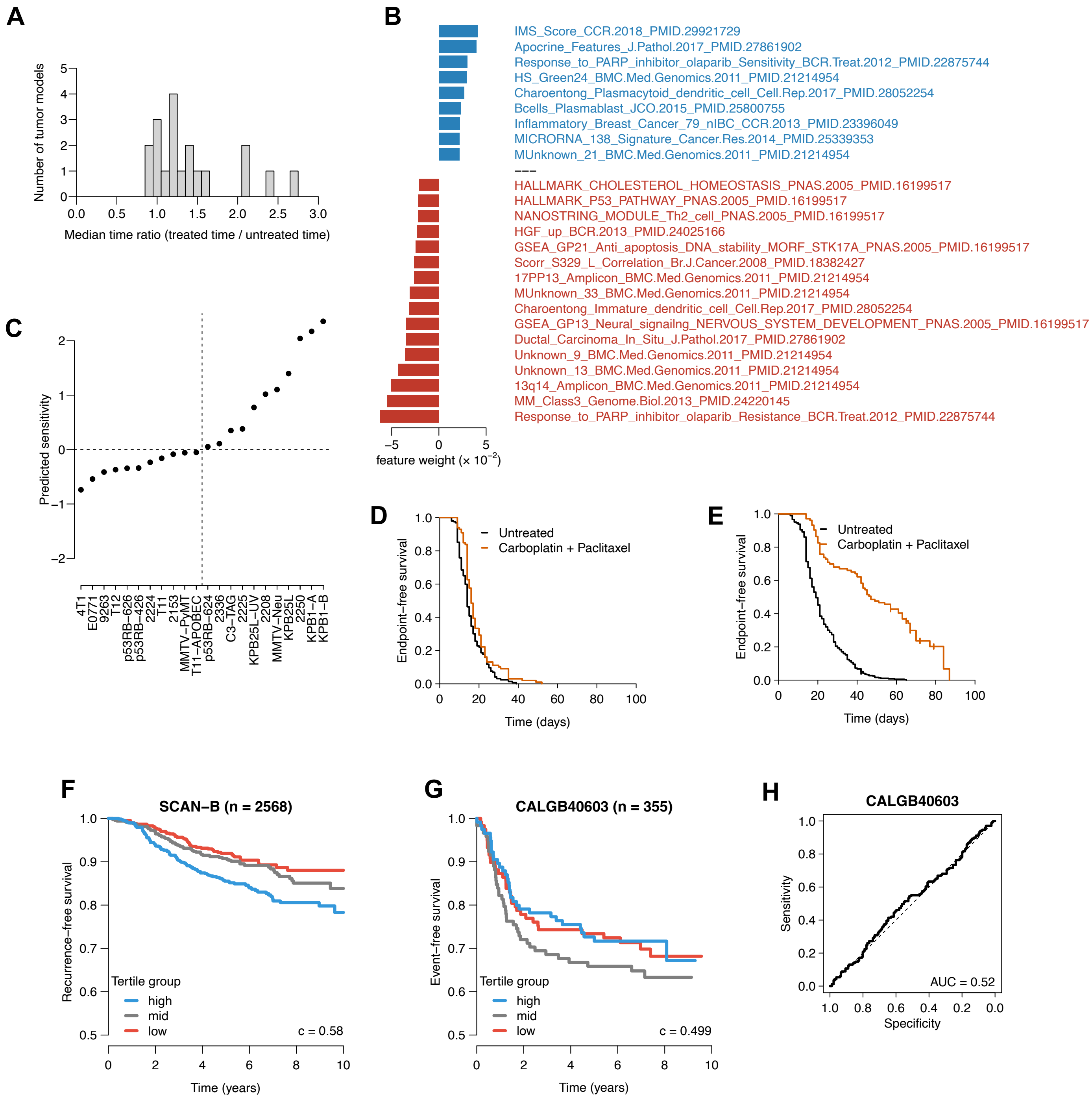
